# Mitochondria-encoded genes contribute to the evolution of heat and cold tolerance among *Saccharomyces* species

**DOI:** 10.1101/390500

**Authors:** Xueying C. Li, David Peris, Chris Todd Hittinger, Elaine A. Sia, Justin C. Fay

**Affiliations:** Molecular Genetics and Genomics Program, Washington University, St. Louis, MO.; Department of Genetics, Washington University, St. Louis, MO.; Center for Genome Sciences and System Biology, Washington University, St. Louis, MO.; Laboratory of Genetics, DOE Great Lakes Bioenergy Research Center, Genome Center of Wisconsin, Wisconsin Energy Institute, J. F. Crow Institute for the Study of Evolution, University of Wisconsin–Madison, Madison, WI.; Department of Food Biotechnology, Institute of Agrochemistry and Food Technology (IATA), CSIC, Paterna, Valencia, Spain.; Department of Biology, University of Rochester, Rochester, NY.

**Author notes:** Corresponding author: Justin C. Fay.

**Keywords:** *Saccharomyces*, non-complementation screen, evolutionary genetics, mitochondria, thermotolerance, cryotolerance

## Abstract

Over time, species evolve substantial phenotype differences. Yet, genetic analysis of these traits is limited by reproductive barriers to those phenotypes that distinguish closely related species. Here, we conduct a genome-wide non-complementation screen to identify genes that contribute to a major difference in thermal growth profile between two *Saccharomyces* species. *S. cerevisiae* is capable of growing at temperatures exceeding 40°C, whereas *S. uvarum* cannot grow above 33°C but outperforms *S. cerevisiae* at 4°C. The screen revealed only a single nuclear-encoded gene with a modest contribution to heat tolerance, but a large effect of the species’ mitochondrial DNA (mitotype). Furthermore, we found that, while the *S. cerevisiae* mitotype confers heat tolerance, the *S. uvarum* mitotype confers cold tolerance. Recombinant mitotypes indicate multiple genes contribute to thermal divergence. Mitochondrial allele replacements showed that divergence in the coding sequence of *COX1* has a moderate effect on both heat and cold tolerance, but it does not explain the entire difference between the two mitochondrial genomes. Our results highlight a polygenic architecture for interspecific phenotypic divergence and point to the mitochondrial genome as an evolutionary hotspot for not only reproductive incompatibilities, but also thermal divergence in yeast.

## Introduction

Species are notably adapted to survive and reproduce in diverse environments. Many of the characteristics that enable them to do so emerged in distant history, and there remains considerable uncertainty as to whether evolution occurred through accumulation of numerous small-effect changes (“micromutationism”) or often involves “major genes” of large effect (Orr and Coyne 1992; Orr 2001). Empirically, there is mixed evidence for the two models (reviewed in Orr 2001; Hoekstra and Coyne 2007; Stern and Orgogozo 2008), but many studies were either limited to identifying genes of large effect or did not resolve major effect loci to individual genes. Notably, in some instances, genes of large effect were dissected to multiple changes of small-effect (McGregor et al. 2007; Frankel et al. 2011; Engle and Fay 2012), providing support for the notion of evolutionary hotspots or bottleneck genes (Stern and Orgogozo 2009). Thus, the resolution of interspecific genetic analysis has direct bearing on understanding the genetic basis of evolutionary change.

Reproductive barriers between species, including hybrid sterility and inviability, present a major challenge to mapping interspecific phenotype differences. While quantitative trait mapping has been successfully applied to closely related, interfertile species (Doebley and Stec 1993; True et al. 1997; Colosimo et al. 2004; Joron et al. 2006; Fishman et al. 2014), the cumulative results of such studies may not be representative of genetic differences between more distantly related species. Characters that distinguish sibling species and domesticated organisms are often related to sexual reproduction (Eberhard 1985) and food production (Purugganan and Fuller 2009; Larson and Fuller 2014) where strong selection and rapid evolution may result in a preponderance of major genes.

Temperature tolerance is a trait that is critical to microbes, plants, and animals without a means of temperature regulation, and it is also variable across species. While the *Saccharomyces* yeast species share their preference for fermentative metabolism with many other yeast species (Hagman et al. 2013; Dashko et al. 2014), they differ dramatically in their thermal growth profile (Gonçalves et al. 2011; Salvadó et al. 2011). The sibling species *S. cerevisiae* and *S. paradoxus* are relatively heat tolerant, capable of growing at temperatures of 37-42°C, while the more distantly related *S. kudriavzevii* and *S. uvarum* are more cold tolerant and only capable of growing at temperatures up to 34-35°C (Gonçalves et al. 2011, Salvadó et al. 2011). Previous studies in yeasts have implicated a small number of genes involved in temperature divergence (Gonçalves et al. 2011; Paget et al. 2014). However, every gene product has the potential to be thermolabile, and only a single systematic screen has been conducted (Weiss et al. 2018), which reported that multiple genes contribute to thermal differences between *S. cerevisiae* and *S. paradoxus*, two species with modest differences in heat tolerance.

In the present study, we examined the genetic basis of thermal divergence between *S. cerevisiae* and *S. uvarum*, two species that are more different at synonymous sites than human and mouse (Waterston et al. 2002; Kawahara and Imanishi 2007). These two species are capable of forming hybrids, but the hybrids are sterile owing to both mitochondrial-nuclear incompatibilities (Lee et al. 2008; Chou et al. 2010) and defects in recombination due to high levels of sequence divergence (Hunter et al. 1996; Liti et al. 2006). Of relevance, mitochondrial genome variation has been associated with temperature-dependent growth differences among *S. cerevisiae* strains (Paliwal et al. 2014; Wolters et al. 2018) and *S. paradoxus* populations (Leducq et al. 2017).

To identify genes involved in the evolution of thermal growth differences, we screened 4,792 non-essential genes for non-complementation. While no genes of large effect were recovered, we found that mitochondrial DNA (mtDNA) plays a remarkable role in divergence of both heat and cold tolerance across the *Saccharomyces* species and that multiple mitochondria-encoded genes are involved, including *COX1*, previously shown to be involved in mitochondrial-nuclear interspecific incompatibilities (Herbert et al. 1992; Chou et al. 2010).

## Results

### A non-complementation screen for thermosensitive alleles reveals mitochondrial effects

In *S. cerevisiae* and *S. uvarum* hybrids, thermotolerance alleles are dominant (Fig. 1A). Thus, deletion of *S. cerevisiae* thermotolerance alleles should weaken thermotolerance through non-complementation. We screened 4,792 non-essential genes in the yeast deletion collection for such thermotolerance genes by mating both the *MAT***a** (BY4741) and *MAT*α (BY4742) deletion collection to *S. uvarum* and growing them at high temperature (37°C). For comparison, we also screened the resulting hemizygote collections for two other traits where the *S. cerevisiae* phenotype is dominant in the hybrid (Fig. 1A): copper resistance (0.5 mM copper sulfate) and ethanol resistance (10% ethanol at 30°C). We found 80, 13, and 2 hemizygotes that exhibited reduced resistance to heat, copper, and ethanol, respectively, in both the BY4741 and BY4742 hemizygote collections (Fig. 1B). In our initial assessment of these genes, we validated a copper-binding transcription factor, *CUP2* (Buchman et al. 1989), for copper resistance through reciprocal hemizygosity analysis (Fig. S1).

**Fig. 1.**
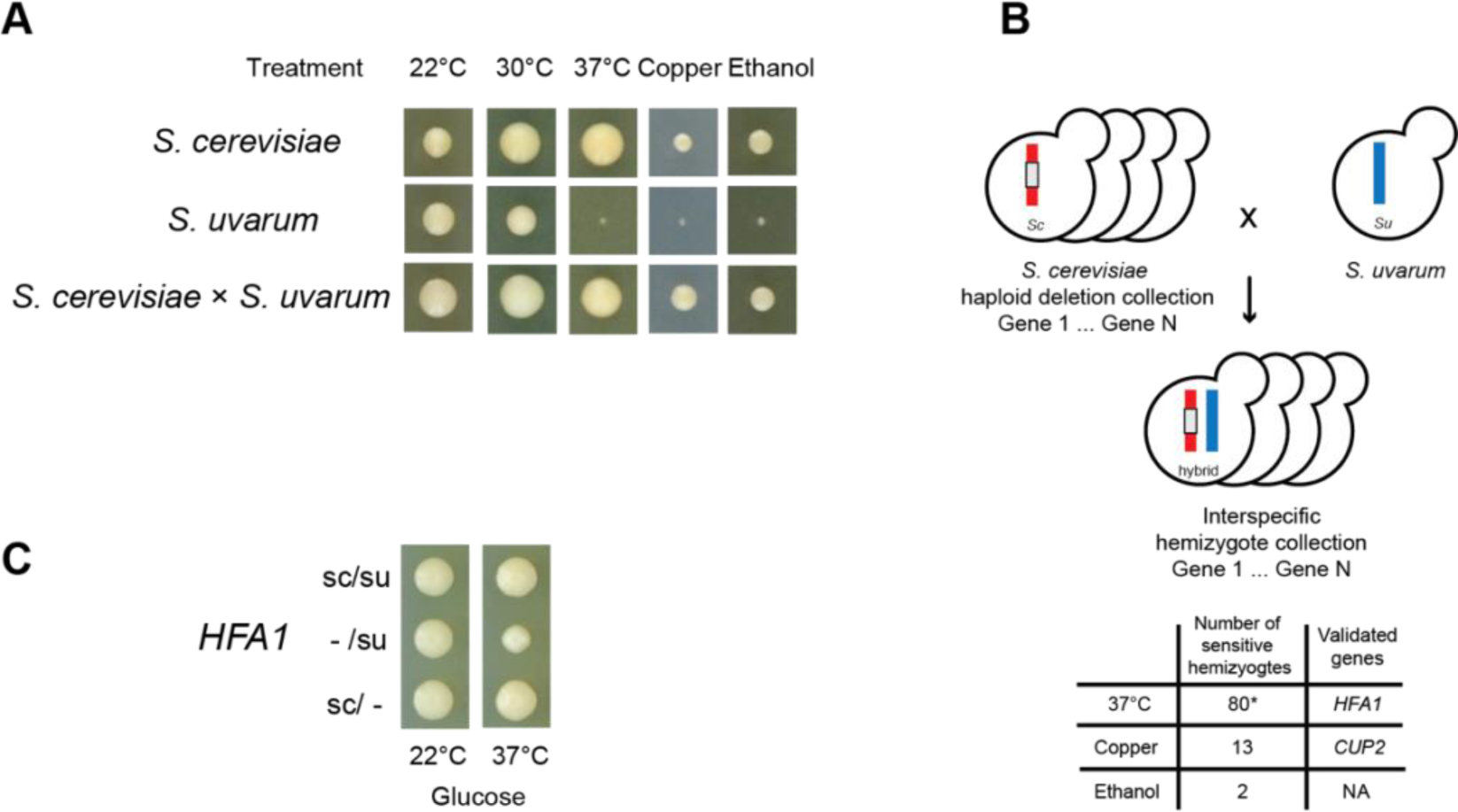
Non-complementation screen for genes underlying phenotypic divergence between *S. cerevisiae* and *S. uvarum*. (**A**) *S. cerevisiae* and *S. uvarum* differ in heat (37°C), copper (0.5mM, 22°C), and ethanol (10%, 30°C) tolerance. The resistant *S. cerevisiae* alleles are dominant, shown by the hybrid (*S. cerevisiae* × *S. uvarum*) compared to *S. cerevisiae* (diploid, S288C background) and *S. uvarum* (diploid, CBS7001 background). Growth is after 3 days. (**B**) *S. cerevisiae* haploid deletion collection was crossed to *S. uvarum* to construct an interspecies hemizygote collection. The number of non-complementing genes is shown for each phenotype; the asterisk indicates the number includes strains carrying *S. uvarum* mtDNA. (**C**) *HFA1* hemizygote with only an *S. cerevisiae* allele (sc/-) shows better 37°C growth than one with only an *S. uvarum* allele (-/su). Growth is after 5 days. See Fig. S1B for quantification.

Nearly all the heat-sensitive hemizygotes (77/80) were from crosses between respiration-deficient (“petite”) *S. cerevisiae* deletion strains and *S. uvarum*. Petites form small colonies on fermentable carbon sources (e.g. glucose) and do not grow on non-fermentable carbon sources (e.g. glycerol). The majority of the petite strains that we identified carried knockouts of nuclear-encoded genes required for mitochondrial function (i.e. “nuclear petites”). However, the effects of these genes can be confounded with mitochondrial DNA (mtDNA) inheritance, which can be from either species. When hybrids of *S. cerevisiae* and *S. uvarum* are constructed they typically fix *S. cerevisiae* mtDNA (Albertin et al. 2013), which they cannot do if the *S. cerevisiae* mtDNA was lost prior to mating. Indeed, PCR-genotyping showed that many of the heat-sensitive hemizygotes carried *S. uvarum* mtDNA. We confirmed one gene (*HFA1*) by reciprocal hemizygosity analysis (Fig. 1C, Fig. S1) causes a moderate loss of heat tolerance due to the *S. uvarum* allele in the presence of *S. cerevisiae* mtDNA. *HFA1* is required for mitochondrial fatty acid biosynthesis (Hoja et al. 2004), and membrane fatty acid composition and fluidity is thought to be important for thermotolerance (Swan and Watson 1997; Swan and Watson 1999)

The inheritance of *S. uvarum* mtDNA in heat-sensitive hemizygotes suggested that mtDNA, rather than the deletion, could be the cause. To test whether the species’ mtDNA (“mitotype”) affects heat tolerance, we generated diploid hybrids of wild-type *S. cerevisiae* and *S. uvarum* with reciprocal mitotypes and grew them at different temperatures. In comparison to the hybrid with *S. cerevisiae* mitotype, the hybrid with the *S. uvarum* mitotype showed reduced fermentative growth at 37°C compared to 22°C and almost no respiratory growth at 37°C (Fig. 2A).

**Fig. 2.**
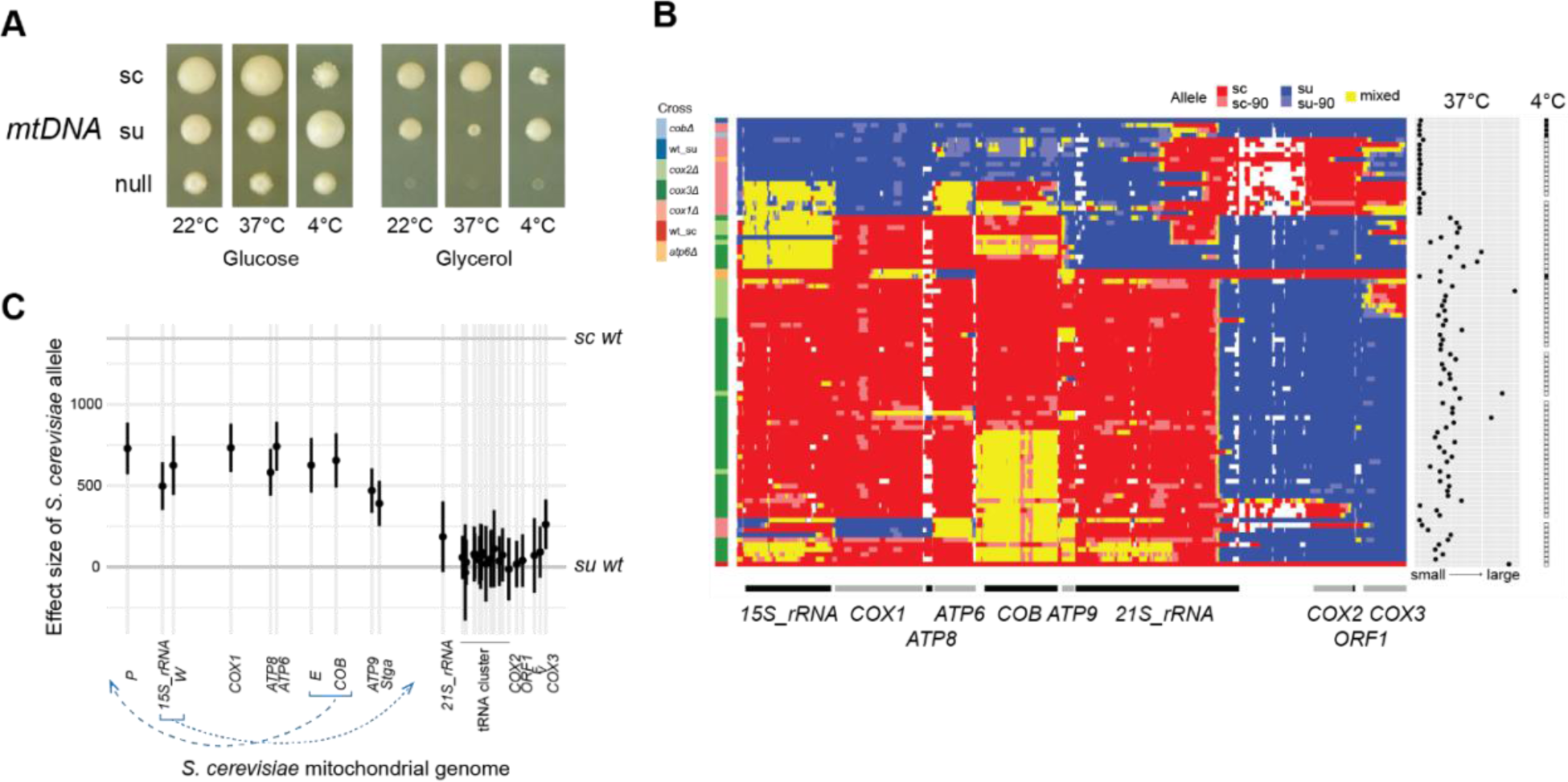
Mitochondria-encoded genes contribute to divergence in heat and cold tolerance. (**A**) *S. cerevisiae* (sc) mtDNA confers heat tolerance to interspecies hybrids, while *S. uvarum* (su) mtDNA confers cold tolerance. (**B**) Recombinant mtDNA genotypes and phenotypes of hybrids. Recombinant strains (rows) derived from mutant crosses (left) are clustered by genotype (middle). Hybrids with wild-type *S. cerevisiae* (wt_sc) and *S. uvarum* mtDNA (wt_su) are at the bottom and the top, respectively. Allele identity is shown for 12.6k single nucleotide markers (sc, *S. cerevisiae*; sc-90, *S. cerevisiae* with a relaxed threshold; su, *S. uvarum*; su-90, *S. uvarum* with a relaxed threshold; mixed, heterozygous or chimeric; white indicates no data). The sites are in the *S. cerevisiae* gene order (bottom), and only sites with orthologous regions are included (site spacing does not reflect mtDNA genome position). 37°C respiratory growth is the average size of non-petite colonies on glycerol plates (right). The presence of 4°C glycerol growth is indicated by solid squares (far right). See Fig. S4A for strain labels. (**C**) Effect size of *S. cerevisiae* alleles on 37°C respiration, with error bars representing 95% confidence intervals. Effect sizes are derived from linear models, and the y-axis is rescaled such that 0 represents the phenotype of wild-type *S. uvarum* mtDNA. The top horizontal line represents the phenotype of wild-type *S. cerevisiae* mtDNA. Selected tRNAs are labeled by their single letter amino acid code. Blue dashed lines indicate genome positions of *S. uvarum* genes compared to *S. cerevisiae*.

*S. uvarum* is not only known to be heat sensitive, but also exhibits enhanced growth at low temperatures relative to *S. cerevisiae* (Gonçalves et al. 2011). We thus tested and found that *S. uvarum* mitotype conferred a growth advantage at 4°C in comparison to *S. cerevisiae* mitotype (Fig. 2A), suggesting a potential trade-off between the evolution of heat and cold tolerance.

To test whether mtDNA-mediated evolution of temperature tolerance is specific to either the *S. cerevisiae* or *S. uvarum* lineages, we generated five additional hybrids with both parental mitotypes using two other *Saccharomyces* species (Fig. S2). In comparison to the 22°C control, we find that both the *S. cerevisiae* and *S. paradoxus* nuclear genome conferred heat tolerance to hybrids with *S. kudriavzevii* and *S. uvarum* (rho° comparison), but the *S. cerevisiae* mitotype conferred heat tolerance in comparison to the *S. paradoxus, S. kudriavzevii*, and *S. uvarum* mitotypes on glucose medium. For cold tolerance we find that the *S. uvarum* mitotype conferred greater cold tolerance relative to the *S. cerevisiae*, *S. paradoxus*, and *S. kudriavzevii* mitotypes. Interestingly, none of the hybrids were as cold tolerant as *S. uvarum* on glycerol. Our results suggest that mtDNA has played an important role in divergence of thermal growth profiles among the *Saccharomyces* species, with heat tolerance evolving primarily on the lineage leading to *S. cerevisiae* and cold tolerance evolving primarily on the lineage leading to *S. uvarum*. A related study has shown these differences have had a direct impact on the domestication of lager-brewing yeast hybrids (Baker et al. 2018)

### Mapping mitochondria-encoded thermotolerant alleles

To identify genes conferring heat tolerance to *S. cerevisiae* mtDNA, we tested whether *S. uvarum* alleles can rescue the respiratory deficiency of *S. cerevisiae* mitochondrial gene knockouts at high temperature. We crossed *S. uvarum* to previously constructed *S. cerevisiae* mitochondrial knockout strains and plated them on glycerol medium at 37°C. Because heteroplasmy is unstable in yeast, this strategy selects for recombinants between the two mitochondrial genomes: *S. uvarum* mtDNA is needed to rescue the *S. cerevisiae* deficiency, and *S. cerevisiae* mtDNA is needed to grow at high temperature (Fig. S3). If the *S. uvarum* gene required for *S. cerevisiae* rescue is temperature sensitive, we expect to see no or small colonies on 37°C glycerol plates. Of the six genes tested, *COX2* and *COX3* deletions were rescued by *S. uvarum* at high temperature, although the colonies were often smaller than the hybrid with wild-type *S. cerevisiae* mtDNA. In contrast, *COX1* and *ATP6* deletions were minimally rescued (Fig. 2B), and *COB* and *ATP8* deletions were not rescued. However, the absence of rescue could also result from a lack of recombination, especially for *COB* because its genomic location has moved between the two species.

Using genome sequencing, we mapped breakpoints in 90 recombinants to determine which *S. cerevisiae* genes are associated with high temperature growth. The recombinants showed hotspots at gene boundaries and within the 21S ribosomal RNA (Fig. 2B). In most cases, the two species’ mtDNA recombine into a circular mitochondrial genome, but sometimes recombination resulted in mitochondrial aneuploidy, particularly for regions where the two species’ mitochondrial genomes are not co-linear (see Fig. S4B for examples). One complication of measuring mtDNA-dependent heat tolerance is the high rate of mtDNA loss, typically 1% in *S. cerevisiae* strains, but much higher in the hybrids and variable among recombinants (Supplementary text, Fig. S5). We thus measured the frequency of petites at 23°C and heat tolerance by the size of single colonies at 37°C on glycerol. We found that the petite frequency was associated with the absence of *S. cerevisiae ORF1* (F-*Sce*III, Peris et al. 2017), a homing endonuclease linked to *COX2* (Fig. S5B & C). For heat tolerance, we found a region including four protein-coding genes (*COX1*, *ATP8*, *ATP6*, and *COB*) with the largest effect (Fig. 2C). The effects associated with these genes are small compared to the total difference between two wild-type mitotypes, suggesting that other regions are required for complete rescue of high temperature growth. Indeed, *S. cerevisiae COX2* and *COX3* showed small but positive effects when the recombinants lacking them were compared to the wild-type *S. cerevisiae* mitotype (Fig. 2B). The differential heat sensitivity is unlikely to be caused by fitness defects since the recombinants grew normally at 23°C (Fig. S4A).

We also found that nearly all mtDNA recombinants did not exhibit 4°C respiratory growth; one strain (S87) derived from the *atp6Δ* cross (Figure 2B) was an exception, but another strain with the same mitochondrial genotype did not grow. The 4°C recombinant phenotypes suggest that cold tolerance might require multiple *S. uvarum* alleles and potentially a different set of genes than those underlying heat tolerance.

### COX1 protein divergence affects both thermotolerance and cryotolerance

Because the recombinant strains did not resolve heat tolerance to a single gene, we tested individual genes by replacing *S. cerevisiae* with *S. uvarum* alleles. We obtained allele replacements for two of the four genes in the region conferring heat tolerance (Fig. 3). For both genes we used intronless alleles to eliminate incompatibilities in splicing (Chou et al. 2010).

**Fig. 3.**
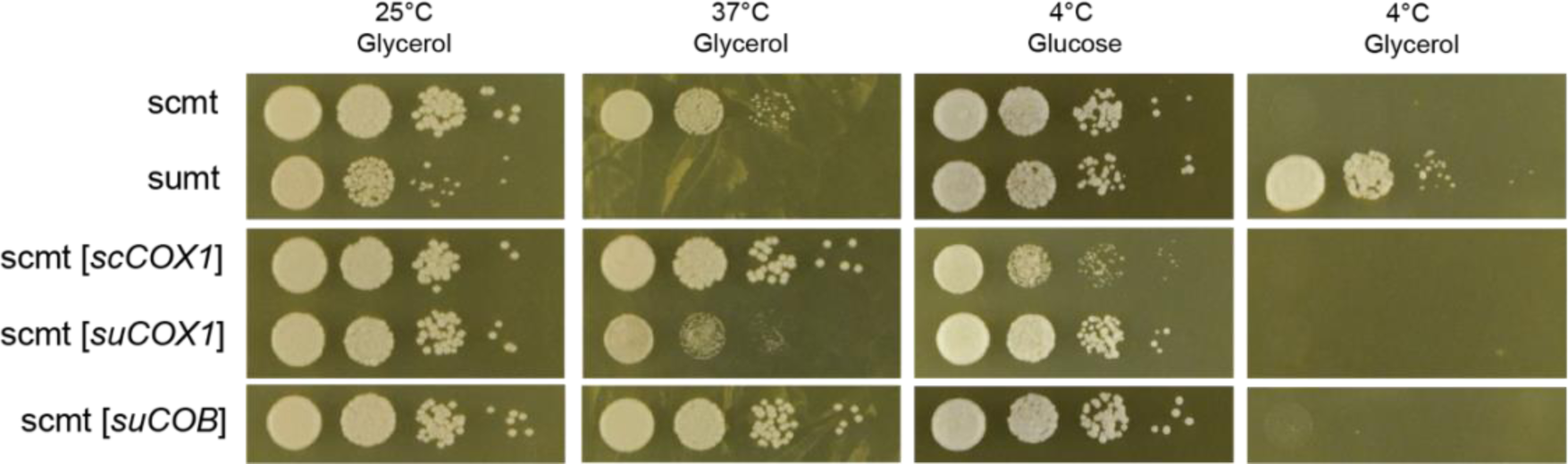
*COX1* coding alleles affect growth at high and low temperature. Hybrids carrying allele replacements and two wild-type controls were plated with 1:10 serial dilution and incubated at indicated temperatures. Growth is after 4 days for 25°C and 37°C, 25 days for 4°C on glucose, 53 days for 4°C on glycerol. sc, *S. cerevisiae*; su, *S. uvarum*; mt, mtDNA. Alleles in the brackets were integrated into their endogenous position in *S. cerevisiae* mtDNA.

We observed a significant difference between *S. cerevisiae* and *S. uvarum COX1* alleles for respiratory growth at 37°C in the hybrid background, with the *S. uvarum* allele being heat sensitive. The effect was not present at room temperature, and the *S. uvarum* allele conferred a growth advantage on glucose at 4°C. Thus, divergence in the *COX1* coding sequence (CDS) affects both heat and cold tolerance. However, *COX1* alleles do not explain the entire difference between the two species’ mitotypes: the strain bearing *S. uvarum COX1* had an intermediate level of heat tolerance and did not confer cold tolerance on glycerol, suggesting that other mitochondrial genes are involved. The moderate effect of the *COX1* alleles is also consistent with the small effect sizes shown by recombinant analysis (Fig. 2C). Surprisingly, the *COX1* allele difference is only seen in the hybrid and not in a diploid *S. cerevisiae* background (Fig. S7), suggesting that the allele difference in hybrid depends on a dominant interaction with the *S. uvarum* nuclear genome.

The *S. uvarum COB* allele replacement rescued respiratory growth at high temperature, demonstrating that protein changes in *COB* do not contribute to heat tolerance. We were unable to generate the *S. cerevisiae* intronless *COB* allele replacement for comparison. Notably, both the intronless *S. cerevisiae COX1* and *S. uvarum COB* allele replacement strains exhibited better growth than wild-type *S. cerevisiae* mtDNA at 37°C (Fig. 3), implying a dominant-negative role of these introns in the hybrid at high temperature.

## Discussion

In *Saccharomyces* species, the mitochondrial genome is not essential for viability, is unusually large for an eukaryote (~86 kb), and is quite variable in intron content (Wolters et al. 2015; Wu et al. 2015). While the mitochondrial genome can recombine and introgress between species (Leducq et al. 2017; Peris et al. 2017), it also contributes to reproductive isolation through incompatibilities with the nuclear genome (Lee et al. 2008; Chou et al. 2010; Spirek et al. 2015). Our results show that the mitochondrial genome also makes a significant contribution to one of the most distinct phenotypic differences among the *Saccharomyces* species: their thermal growth profile. Below, we discuss the implications of our results in relationship to the genetic architecture of species’ phenotypic differences, the role of cyto-nuclear interactions in phenotypic evolution and reproductive isolation, and mitochondria as an evolutionary hotspot for *Saccharomyces* speciation and adaptation.

### Genetic architecture of interspecies differences in thermotolerance

Crosses between closely related, inter-fertile species have shown that phenotypic divergence can be caused by a few loci of large effect, many loci of small effect or a mixture of the two, e.g. (True et al. 1997; Orr 2001; Fishman et al. 2002; Albert et al. 2008). In this study, we carried out a genome-wide non-complementation screen between two distantly related yeast species, with rates of divergence at synonymous sites (1.085, Kawahara and Imanishi 2007) greater than that observed between human and mouse (0.602, Waterston et al. 2002). Out of 4,792 non-essential genes in our study, we found only one gene (*HFA1*) that showed a moderate effect on heat tolerance regardless of the mtDNA effect (Fig. 1C). Of relevance, 178 *S. cerevisiae* deletions are sensitive to 37°C (Auesukaree et al. 2009); a rate comparable to a subsample we examined in this study (78/2251). We can thus conclude that the vast majority of the *S. uvarum* alleles tested exhibited no detectable loss of function at a temperature they do not experience in their native genome. However, our non-complementation screen had some limitations. We did not test essential genes and could not detect genes whose effects were masked by mtDNA inheritance or epistasis, which could occur due to the hybrid carrying an otherwise complete complement of both nuclear genomes.

Although our screen led us to discover a pronounced temperature dependent effect of mtDNA on respiratory growth and a more subtle effect on fermentative growth, the mtDNA effect explains only a small portion of the large difference in heat tolerance between the two species. The *S. cerevisiae* × *S. uvarum* hybrid without mtDNA grows at both 37°C and 4°C on glucose (Fig. 2B), indicating that the nuclear genomes carry dominant factors that remain to be identified.

Despite the small number of genes in the mitochondrial genome, our results show multiple genes within mitochondrial genome influence heat tolerance. In addition to large effect of the *COX1-COB* region, recombinants that inherited *S. uvarum COX2* and/or *COX3* are considerably more heat sensitive than a hybrid with a complete *S. cerevisiae* mtDNA genome. Furthermore, while the *COX1*-linked region showed the largest effect, the *COX1* CDS does not explain the entire difference between two species’ mitotypes. Although we ruled out protein-coding changes in *COB*, changes in the other protein-coding sequences and in gene expression remain to be tested.

The cause of mtDNA-mediated differences in cryotolerance is more opaque. At 4°C, only one recombinant with a significant fraction of *S. cerevisiae* mtDNA grew better than hybrids with an *S. cerevisiae* mitotype, suggesting that multiple *S. uvarum* alleles are required for cold tolerance. Although we showed that *S. uvarum COX1* increased cold tolerance on glucose, the effect is not seen on glycerol, suggesting its effect on respiration might depend on the presence of other *S. uvarum* mitochondrial alleles. However, because the recombinants were all isolated at 37°C, it is possible that they all share some other genetic element or change that facilitates heat tolerance but inhibits 4°C growth.

### Cyto-nuclear interactions in Saccharomyces evolution

In addition to mitochondria-encoded genes, approximately 1,000 nuclear genes function in the mitochondria, many of which are involved in expression and regulation of mitochondrial genes and formation of the multi-subunit cytochrome b and c complexes (Vögtle et al. 2017). Among *Saccharomyces* species, multiple cyto-nuclear incompatibilities have been shown to contribute to reproductive isolation. *S. uvarum AEP2* cannot regulate the translation of *S. cerevisiae ATP9* mRNA (Lee et al. 2008), while *S. cerevisiae MRS1* cannot splice introns of *S. paradoxus* and *S. uvarum COX1* (Chou et al. 2010). Additionally, the *S. uvarum* RNA binding protein *CCM1* has reduced affinity for the *S. cerevisiae* 15s rRNA (Jhuang et al. 2017). While these incompatibilities affect the construction of cybrids, where mtDNA from different species was introduced into *S. cerevisiae* (Spirek et al. 2015), the phenotypic consequences besides loss of respiration is not known.

Our results show that the mitochondrial genomes of *Saccharomyces* species influence both heat and cold tolerance and provide multiple lines of evidence for the role of cyto-nuclear interactions. First, the temperature effects of species’ mitotypes interact with nuclear background (Fig. S1). While *S. cerevisiae* hybrids without mtDNA (rho°) grow similarly on glucose medium, *S. cerevisiae* mtDNA confers different levels of heat tolerance in hybrids with *S. paradoxus*, *S. uvarum*, and *S. kudriavzevii*, the later of which only grows slightly better than the rho° hybrid.

We also observed interactions between the *COX1* allele replacements and their nuclear background. *COX1* showed allele differences at high and low temperatures in the hybrid but not in *S. cerevisiae*. This difference can be explained by a species-specific dominant interaction, as might occur when there are hybrid protein complexes (Piatkowska et al. 2013). In this scenario, *S. uvarum COX1* can function with interacting *S. cerevisiae* proteins at high temperature but exhibits a loss of function when interacting with temperature sensitive *S. uvarum* nuclear factors that are dominant to their *S. cerevisiae* orthologs. The nuclear factor is unlikely to be the previously reported intron-splicing factor *MRS1* because our *COX1* alleles are intronless.

However, introns might affect temperature sensitivity. The intronless *S. cerevisiae COX1* and *S. uvarum COB* alleles showed better respiratory growth at 37°C than wild-type *S. cerevisiae* mtDNA, suggesting a dominant negative role of introns in the hybrid. In *Saccharomyces*, the number and presence of mitochondrial introns is variable between species (Sulo et al. 2017). This contrasts with high conservation of mitochondrial protein coding sequences, which show over 90% sequence identity between *S. cerevisiae* and *S. uvarum*, much higher than the 80% average of nuclear-encoded genes (Kellis et al. 2003). The rapid evolution of introns might require co-evolution of splicing factors, such as *COX1* and *MRS1*. The wild-type hybrid with *S. cerevisiae* mtDNA might be under burden of intron splicing at high temperature caused by dominant negative *S. uvarum* splicing factors. Nevertheless, many introns self-splice and/or encode maturases or homing-endonucleases, which could be temperature sensitive in a nuclear-independent manner.

There is no clear indication that previously reported incompatibilities contribute to the mtDNA temperature phenotypes. The reported cyto-nuclear incompatibilities are recessive, and thus should not contribute to the hybrid phenotypes. For example, although the *S. cerevisiae MRS1* is incompatible with *S. uvarum COX1*, the latter can be correctly spliced by *S. uvarum MRS1* in the diploid hybrid, at least at permissive temperatures. One possibility is that *S. uvarum MRS1* is heat sensitive, which would explain the heat sensitivity of the *S. uvarum* mitotype because neither the *S. cerevisiae* nor *S. uvarum MRS1* would splice *S. uvarum COX1* at high temperature. Heat sensitivity of *S. uvarum MRS1* was tested in our non-complementation screen, but the result was inconclusive. The *S. cerevisiae MRS1* deletion was complemented by the *S. uvarum* allele in the *MAT***a** (BY4741) cross; but its effect was masked by mtDNA inheritance in the *MAT***α** (BY4742) cross. In this regard it is worth noting that *S. cerevisiae* chromosome 9, which carries *MRS1*, is duplicated in three of the recombinant strains; in two cases, these strains show increased 37°C growth compared to similar genotypes (Table S1).

### Mitochondrial DNA and yeast evolution

It has been proposed that mtDNA plays a disproportionate role in Dobzhansky-Muller incompatibilities. Although it is a small genome, it heavily interacts with nuclear genes and has a high nucleotide substitution rate, leading to co-evolution of the mitochondrial and nuclear genomes and multiple interspecific incompatibilities (Burton and Barreto 2012). Has adaptation played a role in driving these incompatibilities? Although no direct links are proven, evolution of the mitochondrial genome and mito-nuclear epistasis has been linked to multiple phenotypes (Solieri et al. 2008; Albertin et al. 2013; Picazo et al. 2014), including 37°C growth (Paliwal et al. 2014, Wolters et al. 2018, Leducq et al. 2017), and deficiencies in mitochondrial DNA cause heat sensitivity (Zubko and Zubko 2014). Here, we show that mtDNA is important for evolution of heat and cold tolerance in distantly related species, caused by the accumulation of multiple small-to-medium effect changes and potentially mito-nuclear epistasis. Taken together, the present and prior findings point to mtDNA as an evolutionary hotspot for yeast speciation and adaptation.

## Materials and Methods

### Strains, growth conditions, and genetic manipulations

*S. uvarum* strains used in this study are derivatives of YJF1449 (*MAT***a** *hoΔ*::*NatMX*) and YJF1450 (*MAT*α* hoΔ*::*NatMX*) in the CBS7001 background (Scannell et al. 2011). *trp1 S. uvarum* strains YJF2600 and YJF2601 were constructed by replacing *TRP1* with *hphMX4* in YJF1449 and YJF1450, respectively. *S. cerevisiae* strain YJF173 (*MAT***a** *ho ura3-52*) in the S288C background was used in reciprocal hemizygosity tests. Strains in other backgrounds will be noted below.

*S. cerevisiae* was maintained on YPD (1% yeast extract, 2% peptone, 2% dextrose) at 30°C; *S. uvarum* and *S. cerevisiae* × *S. uvarum* hybrids were maintained on YPD at room temperature. Strains were also grown on complete medium (CM, 0.3% yeast nitrogen base with amino acids, 0.5% ammonium sulfate, 2% dextrose), or dropout medium (CM-xxx, 0.13% dropout powder, 0.17% yeast nitrogen base, 0.5% ammonium sulfate, 2% dextrose) where xxx represents the missing amino acids when appropriate. SDPSer medium (synthetic dextrose proline D-serine, 2% dextrose, 0.17% yeast nitrogen base without ammonium sulphate or amino acids, 5 mg/ml L-proline, 2 mg/ml D-serine) was used to select for *dsdAMX4* (Vorachek-Warren and McCusker 2004). Antibiotics were added to media when selecting for *KanMX*, *NatMX*, and *hphMX*. YPGly medium (1% yeast extract, 2% peptone, 3% glycerol) was used to examine respiratory growth.

*S. cerevisiae* and *S. uvarum* strains were mated by mixing strains with opposite mating types on YPD at room temperature overnight. Diploid hybrids were obtained by plating the mating mixture to double selection medium and confirmed by mating-type PCR.

Transformations in this study followed standard lithium acetate methods (Gietz et al. 1995). When transforming *S. uvarum* or *S. cerevisiae* × *S. uvarum* hybrid, we used 37°C for heat shock and room temperature for incubation.

Strains lacking mitochondrial DNA (rho^0^) were generated by overnight incubation with shaking in liquid minimal medium (MM, 0.17% yeast nitrogen base without amino acid and ammonium sulfate, 0.5% ammonium sulfate, 2% dextrose) containing 10 mg/ml ethidium bromide. Following incubation, the culture was plated to YPD and YPGly to identify non-respiring colonies.

### Interspecific hemizygote collections

The haploid yeast deletion collections derived from BY4741 (*MAT***a** *his3Δ1 leu2Δ0 met15Δ0 ura3Δ0*) and BY4742 (*MATα his3Δ1 leu2Δ0 lys2Δ0 ura3Δ0*) were arrayed in 384-well format using a Singer ROTOR (Singer Instruments, Watchet UK) and mated to *trp1 S. uvarum* strains. Diploids were selected on CM-trp-his-leu-lys-ura plates. The resulting two interspecific hybrid collections were hemizygous for 4,792 genes.

The hemizygote collections were screened for non-complementation using the following conditions: 1) YPD at room temperature, 30°C, 35°C, and 37°C; 2) CM with 0.5 mM copper sulfate at room temperature; and 3) YPD with 10% ethanol at 30°C. Pictures of plates were taken on the second and fifth day of incubation using a Nikon D3100 camera. Small colonies on day 5 were scored as sensitive strains. For heat, copper, and ethanol stresses, respectively, we found 145, 137, and 26 non-complemented genes from the BY4741 (*MAT***a**) cross and 221, 134, and 19 from the BY4742 (*MAT***α**) cross, resulting in an intersection of 80, 13, and 2 genes (Data S1).

Respiration-deficient strains (petites) were identified by plating the haploid deletion collection strains on YPGly at 30°C. To estimate the rate of temperature-sensitive deletions, we sampled six plates (~2.3k strains) from the haploid deletion collections and assayed their growth on YPD plates at room temperature and 37°C.

### Validation of non-complementing genes

We first repeated the non-complementation test in another strain background. We made deletions of candidate genes (*HFA1* for heat; *TDA1*, *TDA9*, *GGC1*, *TDA4*, *RPL39*, *ADD66*, *YOL075C*, *CUP2*, and *CAJ1* for copper) by *KanMX* in an *S. cerevisiae* strain YJF173 in the same way as the deletion collection, with the exception that the coding region of *HFA1* was defined according to Suomi et al. (2014). The knockout strains were then crossed to an *S. uvarum* rho^0^ strain. Phenotypes of the hemizygotes were assessed at the same conditions as in the screen, and only phenotypes of *HFA1* and *CUP2* were replicated.

Reciprocal hemizygotes were generated for *HFA1* and *CUP2*. Orthologs of *S. cerevisiae HFA1* and *CUP2* were knocked out in *S. uvarum* strain YJF1450 with *KanMX*. The orthologs were defined according to Scannell et al. (2011); for *HFA1*, we included an extra 477 bp upstream of the ATG for the *S. uvarum* allele, based on translation from a non-AUG start codon at position -372 in *S. cerevisiae* (Suomi et al. 2014). The *S. uvarum* deletion strains were then crossed to *S. cerevisiae* (YJF173), and the resulting hemizygotes were genotyped by PCR and found to carry *S. cerevisiae* mtDNA. Phenotypes of the two reciprocal hemizygotes were assessed on the same plate, under the same conditions as in the screen.

### Interspecific hybrids with reciprocal mitotypes

*Saccharomyces* strains used in generating interspecific hybrids with reciprocal mitotypes are as follows: YJF153 (*MAT***a** *hoΔ*::*dsdAMX4*), derived from an *S. cerevisiae* oak tree isolate YPS163; YJF694 (*MATα hoΔ*::*lox-KAN-lox*) and YJF695 (*MAT***a** *hoΔ*::*lox-KAN-lox*), derived from an *S. paradoxus* oak isolate N17; CR85 (*MATα hoΔ*::*hygMX4 ura3*), a wild *S. kudriavzevii* strain isolated from a Spanish *Quercus ilex* bark provided by Amparo Querol (Lopes et al. 2010); YJF1450 (*MATα hoΔ*::*NatMX*), derived from *S. uvarum* CBS7001; YJF2784 (*MAT***a** *hoΔ*::*dsdAMX4*) and YJF2785 (*MATα hoΔ*::*dsdAMX4*), derived from a *S. uvarum* wine isolate BMV58 and provided by Javier Alonso del Real Arias and Amparo Querol.

Interspecific hybrids with reciprocal mitotypes were generated by crossing a rho^+^ strain from one species to a rho^0^ strain from another species. mtDNA was confirmed by PCR using primers targeting the tRNA clusters in mtDNA (forward 5’-CCATGTTCAAATCATGGAGAGA-3’, reverse 5’-CGAACTCGCATTCAATGTTTGG-3’; 95°C 2min; 95°C 30s, 50°C 30s, 72°C 30s for 30 cycles; 72°C 5min). The expected product sizes are 167 bp for *S. cerevisiae*, 131 bp for *S. paradoxus*, 218 bp for *S. kudriavzevii*, and 100 bp for *S. uvarum*.

### Crosses with mitochondrial knockouts

*S. uvarum* strain YJF2600 (*MAT***a** *hoΔ::NatMX trp1Δ::hphMX4*) and YJF2601 (*MATα hoΔ::NatMX trp1Δ::hphMX4*) were crossed to the following *S. cerevisiae* mitochondrial knockout strains: XPM10b (D273-10B background, *MAT***α** *arg8::hisG leu2-3,112 lys2 ura3-52 cox1Δ::ARG8m rho+*) (Perez-Martinez et al. 2003); HMD22 (D273-10B background, *MAT***a** *ura3-52 leu2-3,112 lys2 his3-deltaHinDIII arg8::hisG rho+ cox2Δ::ARG8m*) (Bonnefoy and Fox 2000); DFS168 (D273-10B background, *MAT***a** *leu2-3,112 his4-519 ura3Δ arg8Δ::URA3, cox3Δ::ARG8m, rho+*) (Steele et al. 1996); CAB59 (D273-10B background, *MAT***α** *arg8::hisG leu2-3,112 lys2 ura3-52 cobΔ::ARG8m COX1* minus at least one intron, *rho+*) (Ding et al. 2009); MR6 ΔATP6 (*MAT***α** *ade2-1 his3-11,15 leu2-3,112 trp1-1 ura3-1 CAN1 arg8::HIS3, rho+ atp6Δ::ARG8m*) (Rak et al. 2007); MR6 ΔATP8 (*MAT***α** *ade2-1 his3-11,15 leu2-3,112 trp1-1 CAN1 ura3-1 arg8::HIS3, rho+ atp8Δ::ARG8m*) (Rak and Tzagoloff 2009). *S. cerevisiae* strains with wild-type mtDNA, PJD1 (D273-10B background, *MAT***α** *ura3-52 ade2-101 leu2-3, 112 ade3-24 rho+*) and MR6 (*MAT***a** *ade2-1 his3-11,15 leu2-3,112 trp1-1 ura3-1 CAN1 arg8::HIS3*), were crossed in parallel as control. The D273-10B wild-type, *cox1Δ*, *cox2Δ*, *cox3Δ*, and *cobΔ* strains were provided by Tom D. Fox; the MR6 wild-type, *atp6Δ* and *atp8Δ* strains were provided by Alexander Tzagoloff.

*MAT***a** and *MAT***α** strains were mixed on YPD and incubated at room temperature overnight. The mating mixtures were either replica-plated (initial trial) or resuspended in sterile water and plated (second trial) onto YPGly. The YPGly plates were incubated at 37°C for 7-10 days to select for 37°C-respiring recombinants. The mating mixtures of *cox2Δ* and *cox3Δ* crosses were also plated to CM-trp-his-leu-lys-ura at room temperature to select for diploid hybrids, which allowed us to estimate the recombination rate to be around 0.05-0.1%. 37°C-respiring colonies were picked and streaked on YPD at room temperature for single colonies. For the initial trial, the 37°C-respiring cells were streaked on YPD twice. For the *cox1Δ* and *atp6Δ* crosses, the plates were left at room temperature for 3 days after 7 days at 37°C incubation and colonies growing from the recovery period were also picked and streaked. As a result, 3+12, 4+48, 3+25, 2+3, 0+7, and 0+1 strains (initial trial + second trial) from the *cox2Δ, cox3Δ, cox1Δ, cobΔ, atp6Δ*, and wild-type D273-10B control crosses, respectively, were generated. The total of 102 strains were subjected to whole genome sequencing and phenotyping.

### Spontaneous mitochondrial recombinants

*S. cerevisiae* (YJF153, *MAT***a** *hoΔ*::*dsdAMX4*, YPS163 derivative) and *S. uvarum* (YJF1450, *MAT***α** *hoΔ*::*NatMX*, CBS7001 derivative) were mated and streaked onto SDPSer + clonNAT medium to select for diploid hybrids. 384 colonies on the double selection plates were picked and arrayed onto one YPD agar plate and subsequently pinned to YPD and YPGly and incubated at room temperature, 37°C and 4°C. Colony sizes on each plate were scored both manually and quantitatively using ImageJ (Rasband 1997). Strains with recombinant-like temperature phenotypes (r114, r194, r262, r334, r347 and b2), along with two control strains (r21, r23) with typical phenotypes for *S. cerevisiae* and *S. uvarum* mitotypes, respectively, were subjected to whole genome sequencing and phenotyping.

### DNA extraction, library preparation, and sequencing

For the unselected putative recombinants and their controls (r21, r23, b2, r334, r114, r194, r262, and r347), DNA was extracted using an mtDNA-enriching protocol (see below). For other strains sequenced in this study, genomic DNA was extracted from 22°C YPD overnight cultures inoculated with cells pre-grown on YPGly plates (ZR Fungal/Bacterial DNA MicroPrep kit, Zymo Research).

mtDNA was enriched following a protocol adapted from Fritsch et al. (2014) and Wolters et al. (2015). 50ml YPEG (1% yeast extract, 2% peptone, 2% ethanol, 2% glycerol) medium was inoculated with overnight YPD starter cultures, shaken at 300rpm at 23°C. The culture was collected at late-log phase (3,000g for 1 min) and the cell pellet was washed twice in 1ml sterile distilled water. The cells were then washed in buffer (1.2M Sorbitol, 50mM Tris pH 7.4, 50mM EDTA, 2% beta-mercaptoethanol) and centrifuged at 14,000 rpm for 3 minutes. The cell pellet was weighed, resuspended in Solution A (0.5M Sorbitol, 50mM Tris pH 7.4, 10mM EDTA, 2% beta-mercaptoethanol, 7ml/g wet weight cells) containing 0.2mg/ml Zymolyase (Zymo Research), and incubated at 37°C, 100 rpm for 45min for osmotic lysis. The suspension was then centrifuged at 4,000rpm for 10min. The supernatant was decanted to a new tube and centrifuged at 14,000 rpm for 15min, to get the crude mitochondrial pellet. The pellet was then incubated in DNase treatment solution [0.3M Sucrose, 5mM MgCl_2_, 50mM Tris-HCl pH 8.0, 10mM CaCl_2_, 100U/ml RQ1 DNase (Promega), 500ul/g initial wet weight] at 37°C, 100 rpm for 30min to remove nuclear DNA. 0.5M EDTA (pH 8.0) was added to a final concentration of 0.2M to stop the reaction. The mitochondrial pellet was then washed three times by repeated cycles of centrifugation at 15,000 rpm for 10min and resuspension in 1ml Solution A to remove DNase, and then resuspended in 400ul Solution B (100mM NaCl, 10mM EDTA, 50mM Tris pH 8) and incubated at room temperature for 30min for lysis. mtDNA was isolated from the solution by phenol-chloroform extraction and ethanol precipitation, followed by a clean-up with DNA clean and concentrator -5 kit (Zymo Research). Alternatively, two samples (r21 and r262) were extracted with ZR Fungal/Bacterial DNA MicroPrep Kit (Zymo Research) by adding the Fungal/Bacterial DNA binding buffer to the lysed mitochondrial fraction and following the rest of the manufacturer protocol. The yield was typically 10-20ng/g wet weight cells and provided 10- to 100-fold enrichment of mitochondrial reads.

Paired-end libraries were prepared with Nextera DNA Library Preparation Kit (Illumina) with a modified protocol. Briefly, 3-5 ng DNA was used for each sample and the tagmentation reaction was performed at a ratio of 0.25ul tagmentation enzyme/ng DNA. The tagmented DNA was amplified by KAPA HiFi DNA polymerase for 13 cycles (72°C 3min; 98°C 5min; 98°C 10s, 63°C 30s, 72°C 30s for 13 cycles; 72°C 5min). The PCR reaction was then purified with AMPure Beads. Paired-end 2x150 Illumina sequencing was performed on a MiniSeq by the DNA Sequencing Innovation Lab in the Center for Genome Sciences and System Biology at Washington University. 96 recombinants generated in the second trial of the mitochondrial mutant crosses were subsequently re-sequenced on a NextSeq 500 at Duke Center for Genomic and Computational Biology for deeper coverage. The NextSeq reads and MiniSeq reads were combined in the analysis. All reads were deposited at the Sequence Read Archive under accession no. SRP155764.

### Mitochondrial genome assembly

The *S. uvarum* mitochondrial genome was assembled from high-coverage sequencing of r23. Before assembly, we confirmed that it carried a non-recombinant *S. uvarum* mitochondrial genome by mapping the reads to CBS380 (Okuno et al. 2016), an *S. eubayanus* × *S. uvarum* × *S. cerevisiae* hybrid that inherited the mitochondria from *S. uvarum*. To assemble the mitochondrial genome, reads were first cleaned with trimmomatic (Bolger et al. 2014) to remove adapters. They were then assembled using SPAdes assembler (Bankevich et al. 2012), included in the wrapper iWGS (Zhou et al. 2016), to produce contigs. Contigs were scaffolded to produce the final assembly through comparison with the output assembly of MITObim (Hahn et al. 2013). The assembly was annotated with MFannot Tool (http://megasun.bch.umontreal.ca/cgi-bin/mfannot/mfannotInterface.pl); *ORF1* (*F-SceIII*) annotation was added manually using Geneious R6 (Kearse et al. 2012). The assembled r23 mitochondrial genome is 64,682 bp and has a total of 5,874 gapped bases. Most gaps are in the intergenic regions, one gap is in *VAR1*, and 3 small gaps are in the introns of *COB*. The r23 mitochondrial genome is 99% identical to CBS380, based on BLAST results. The assembly was deposited at GenBank under accession no. MH718505.

### Read mapping and allele assignment of recombinants

Illumina reads were mapped to a reference that combined the mitochondrial genomes of *S. cerevisiae* (S288C-R64-2-1) and *S. uvarum* (r23 mitochondria assembled in this study), using end-to-end alignment in Bowtie 2 (Langmead and Salzberg 2012). Duplicated reads and reads with high secondary alignment scores (XS>=AS) or low mapping quality (MQ < 10) were filtered out. Using this method, reads from hybrids with non-recombinant *S. cerevisiae* or *S. uvarum* mtDNA were >99.9% correctly mapped to their reference genomes (49,496/49,504 for *S. cerevisiae*, 161,712/161,714 for *S. uvarum*). To characterize aneuploidy and the ratio of mitochondrial to nuclear reads, the reads were re-mapped to a reference file combining *S. cerevisiae* (S288C-R64-2-1) and *S. uvarum* (Scannell et al. 2011) reference genomes, using the same method. Coverage of nucleotide positions and of chromosomes was generated by *samtools depth* and *samtools idxstats*, respectively.

For data visualization and identification of recombination breakpoints, we assigned allele identity for each nucleotide in orthologous regions in the two reference mitochondrial genomes. The total length of orthologous sequences is 16.5kb (nucmer alignment) and contains mostly coding and tRNA sequences. After removing sites with no coverage in control strains, 12.6k nucleotide positions were subjected to data visualization and allele calling. We called the allele identity of a given nucleotide position based on the ratio of reads that mapped to the *S. cerevisiae* reference allele to the total number of reads that mapped to the two orthologous alleles (rsc=sc/(sc+su)): rsc of 1 (or no lower than the non-recombinant *S. cerevisiae* mtDNA control) was called *S. cerevisiae*, rsc of 0 (or no higher than the non-recombinant *S. uvarum* mtDNA control) was called *S. uvarum*, rsc > 0 and < 1 were called mixed. Sites without coverage of either allele were treated as missing data. A relaxed threshold was used in data visualization to account for noise in read mapping (rsc >0.9 was called *S. cerevisiae*, labeled as “sc-90”; rsc <0.1 was called *S. uvarum*, labeled as “su-90”). Using this method, a total of 90 sequenced strains were confirmed to be recombinants.

For quantifying the effect size of *S. cerevisiae* alleles, we counted the number of reads mapped to each protein-coding gene, tRNA and rRNA by htseq-count. For each gene, we tested the allele effect across 90 recombinants using a linear model: *phenotype* ~ *allele* + *petite*, where *allele* is the ratio of *S. cerevisiae* reads for a given gene and *petite* is the empirically determined petite rates (see below). Because we used the ratio of *S. cerevisiae* reads to represent allele identity, the model does not assume dominance; a heterozygous individual (i.e. read ratio = 0.5) should have an intermediate phenotype. P-values were extracted from the models and adjusted by the false discovery rate (Benjamini & Hochberg method) to correct for multiple comparisons. While the p-value for the *petite* term is significant in some models, its effect was always estimated to be positive. Because high petite rates should lead to small colonies, we do not consider petite rate to significantly contribute to the phenotype. Additionally, aneuploidy (Table S1) and mtDNA copy number variation (Data S2) were present in several recombinants, but the addition of the two variables to the model did not change the effect size and significance of the *allele* term (*phenotype* ~ *allele* + *petite*+ *aneuploidy* + *copy*, where *aneuploidy* is a binary variable indicating presence/absence of chromosomal duplication and *copy* is the ratio of mitochondrial to nuclear reads).

The unselected putative recombinants were sequenced to high coverage, so we generated contigs and assemblies as in “Mitochondrial genome assembly”. The contigs were mapped to *S. cerevisiae* (r21) and *S. uvarum* (r23/CBS380/CBS7001) assemblies in Geneious R6 to identify the breakpoints. For the recombinants of lower quality assemblies (r194, r347, and b2), the contigs were mapped to the best recombinant assembly r114 to improve recombinant construction. Results were confirmed by retaining the Illumina reads from the mitochondrial genome, using both reference mitochondrial genomes as baits in HybPiper (Johnson et al. 2016) and mapping them to the reference mitochondrial genomes using Geneious R6.

### Recombinant phenotypes

Recombinant strains were first grown on YPGly plates to enrich for respiring cells, then in liquid YPD shaken at room temperature overnight. The overnight culture was diluted 1:10^5^, spread on YPD and YPGly plates and incubated at 22°C, 37°C, or 4°C. Pictures of plates were taken on the 5^th^ day for 22°C and 37°C YPD plates, on the 6^th^ day for 22°C and 37°C YPGly plates and on the 68^th^ day for 4°C YPD and YPGly plates. Colony sizes on YPGly plates were acquired by the *Analyze Particles* function in ImageJ (Rasband 1997). Non-single colonies were filtered out both by manually marking problematic colonies during analysis and by roundness threshold (roundness > 0.8 for non-petite colonies). For each strain, sizes of all the non-petite colonies (colony size > 200 units) were averaged; if no cells were respiring at a given condition, the average of all the (micro)colonies was used instead. Petite rates of the overnight cultures were recorded by counting big/small colonies on 22°C YPD and normal/micro colonies on 22°C YPGly plates, and the two values were averaged. Control strains carrying wild-type *S. cerevisiae* or *S. uvarum* mtDNA in the background of D273-10B × CBS7001 were phenotyped in parallel.

Initially the ~90 strains were phenotyped in three batches. We accounted for the batch effect for the 37°C data by picking 3-4 strains from each batch and repeating the phenotyping process on the same day at 37°C. Linear models between old data and new data were generated for each batch separately and were used to adjust for an overall batch effect. The 22°C colony sizes were not adjusted.

### Mitochondrial allele replacement

Mitochondrial transformation was performed as previously described (Bonnefoy and Fox 2001) (Fig. S6). Intronless mitochondrial alleles were synthesized by Biomatik. The alleles were Gibson-assembled into an *ARG8m*-baring pBluescript plasmid, such that the mitochondrial allele is flanked by 69bp and 1113 bp *ARG8m* sequences at its 5’ and 3’ end, respectively (Fig. S6C). Sequences of the assembled plasmid were confirmed by Sanger sequencing.

Strains described in the “Crosses with mitochondrial knockouts” section were used in the mitochondrial transformation. They were first transformed with *P_GAL_-HO* to switch mating types and validated by mating type PCR. In these strains, the target gene was replaced with *ARG8m*, so our constructs carrying the allele of interest can integrate into their endogenous loci by homologous recombination with *ARG8m* (Fig. S6C).

We bombarded the mitochondrial plasmid and pRS315 (CEN plasmid carrying *LEU2*) into *S. cerevisiae* strain DFS160 (*MATα ade2-101 leu2Δ ura3-52 arg8Δ::URA3 kar1-1*, *rho^0^*) using a biolistic PDS-1000/He particle delivery system (Bio-Rad) and selected for Leu+ colonies on MM plates. The colonies were replica-mated to the mitochondrial knockout strains at 30°C for 2 days. The mating mixtures were replica-plated to YPGly plates and incubated at 30°C. YPGly+ colonies were streaked on YPD and mating types were determined by PCR. We also isolated the DFS160-derived parent strains that gave rise to the YPGly+ colonies from the master plates. For *S. cerevisiae COX1* and *COB* alleles, the parent strains were re-mated to the knockout strains for confirmation.

The YPGly+ colonies carry a mitochondrial genome with the allele of interest integrated at their endogenous loci. Because of the *kar1-1* mutation in DFS160, we were able to isolate YPGly+ colonies that are diploid, *MAT***a** haploid, or *MATα* haploid. We crossed the *MAT***a** transformant (D273-10B background) to an *S. uvarum* rho^0^ strain. The hybrid strain and the diploid *S. cerevisiae* strains directly obtained from the mitochondrial transformation were phenotyped at room temperature, 37°C, and 4°C on YPD and YPGly by spot dilution assays. The allele identity of all the phenotyped strains was confirmed by PCR and restriction digest.

## Acknowledgements

We thank Tom D. Fox, Alexander Tzagoloff, Javier Alonso del Real Arias, and Amparo Querol for sharing strains. We thank members of Fay lab for comments and experimental assistance. This work was supported by a National Institutes of Health grant (GM080669) to JCF. Additional support to CTH was provided by the USDA National Institute of Food and Agriculture (Hatch project 1003258), the National Science Foundation (DEB-1253634), and the DOE Great Lakes Bioenergy Research Center (DOE BER Office of Science DE-SC0018409 and DE-FC02-07ER64494 to Timothy J. Donohue). CTH is a Pew Scholar in the Biomedical Sciences and a Vilas Faculty Early Career Investigator, supported by the Pew Charitable Trusts and the Vilas Trust Estate, respectively. DP is a Marie Sklodowska-Curie fellow of the European Union’s Horizon 2020 research and innovation programme, grant agreement No. 747775.

